# Neuronal TDP-43 aggregation drives changes in microglial morphology prior to immunophenotype in amyotrophic lateral sclerosis

**DOI:** 10.1101/2024.10.17.618941

**Authors:** Molly E. V. Swanson, Miran Mrkela, Clinton Turner, Maurice A. Curtis, Richard L. M. Faull, Adam K. Walker, Emma L. Scotter

## Abstract

Microglia are the innate immune cells of the brain with the capacity to react to damage or disease. Microglial reactions can be characterised in post-mortem tissues by assessing their pattern of protein expression, or immunophenotypes, and cell morphologies. We recently demonstrated that microglia have a phagocytic immunophenotype in early-stage ALS but transition to a dysfunctional immunophenotype by end stage, and that these states are driven by TAR DNA-binding protein 43 (TDP-43) aggregation in the human brain. However, it remains unclear how microglial morphologies are changed in ALS. Here we examine the relationship between microglial immunophenotypes and morphologies, and TDP-43 pathology in motor cortex tissue from people with ALS and from a TDP-43-driven ALS mouse model. Post-mortem human brain tissue from 10 control and 10 ALS cases was analysed alongside brain tissue from the bigenic *NEFH*-tTA/*tetO*-hTDP-43ΔNLS (rNLS) mouse model of ALS at distinct disease stages. Sections were immunohistochemically labelled for microglial markers (HLA-DR, CD68, and Iba1) and phosphorylated TDP-43 (pTDP-43). Single-cell microglial HLA-DR, CD68, and Iba1 average intensities, and morphological features (cell body area, process number, total outgrowth, and branch number) were measured using custom image analysis pipelines. In human ALS motor cortex, we identified a significant change in microglial morphologies from ramified to hypertrophic, which was associated with increased Iba1 and CD68 levels. In the rNLS mouse motor cortex, the microglial morphological changed from ramified to hypertrophic and increased Iba1 levels occurred in parallel with pTDP-43 aggregation, prior to increases in CD68 levels. Overall, the evidence presented in this study demonstrates that microglia change their morphologies prior to immunophenotype changes. These morphological changes may prime microglia near neurons with pTDP-43 aggregation for phagocytosis, in turn triggering immunophenotype changes; first, to a phagocytic state then to a dysfunctional one.

## Introduction

Microglia are the innate immune cells of the brain, with functional roles in both health and disease. Microglia have the capacity to respond to damage and disease, performing functions such as phagocytosis, cytokine secretion, migration, and antigen presentation [1–13]. To fulfil and switch between these functions, microglia change their morphologies and protein expression, and such changes have been identified in a range of neurodegenerative diseases including amyotrophic lateral sclerosis (ALS) [14–30]. However, while these microglial changes are likely neuroprotective in early-stage ALS, there is evidence that microglia become dysfunctional and neurotoxic in late-stage disease [31–33].

ALS is the most common form of motor neuron disease, characterised by progressive death of upper and lower motor neurons, which leads to muscle atrophy, disability, and death [34]. Diverse genetic and environmental factors cause or increase the risk of ALS [35,36]. Despite this range of risk factors, 97% of all ALS cases share a common pathological hallmark: intracellular accumulation of TAR DNA-binding protein 43 (TDP-43) aggregates in motor neurons [37]. Although TDP-43 aggregates are unlikely to be the ‘toxic species’ in ALS, they are surrogates for the presence of the aggregation pathway species that *are* thought to be toxic; misfolded monomers and oligomers [38]. While motor neurons are central to ALS pathogenesis and symptomatology, microglial immunophenotypes (pattern of protein expression) and morphologies are also altered in response to TDP-43 aggregation [20,32,39–41]. Indeed, in the human ALS brain, microglia up-regulate markers suggestive of increased phagocytosis and dysfunction [39]. However, there has been limited consideration of microglial morphology changes in conjunction with immunophenotypes in the ALS brain.

Microglial morphologies have been examined in post-mortem studies to infer function and/or dysfunction in the ageing and diseased brain [42–51]. Five distinct morphologies have been described: ramified (small cell bodies and complex processes), hypertrophic (large cell bodies with reduced process complexity), dystrophic (small cell bodies with beaded, discontinuous, or tortuous processes), amoeboid (round cell body with pseudopodia), and rod (elongated appearance with small cell body and retraction of planar processes) [42–44,46,50,52–57]. While analyses of microglial morphology in ALS are limited, immunohistochemical labelling of a cytoskeletal protein expressed by all microglia, Iba1, has been used to identify morphological changes in rodent *SOD1* ALS models [15,17,18,21]. Microglia in control mice and transgenic rodents at early disease stages have small cell bodies with complex processes, suggestive of a ramified ‘resting’ morphology. In contrast, microglia in ALS transgenic mice at later disease stages have larger cell bodies with reduced process complexity, suggestive of a hypertrophic ‘reactive’ morphology [15,17,18,21]. A recent study demonstrated that rod-shaped microglia may play a unique neuroprotective role in TDP-43 mice and human sporadic ALS; however, hypertrophic and dystrophic microglia classically identified in ALS brains were not described [41]. Further investigations into how microglial morphologies change in response to the neuronal TDP-43 aggregation characteristic of more than 97% of ALS cases and how these changes relate to function are needed.

While microglial morphology can be used to infer microglial functional states, microglial functions are not thought to be specific to a given morphology. Numerous microglia classification paradigms that consider microglial morphologies and immunophenotypes have been proposed to describe microglial responses in the human brain, including but not limited to; activated versus resting microglia, M1/M2 pro- and anti-inflammatory microglia, and disease-associated microglia (DAMs) [51]. However, these somewhat binary classifications of microglial states do not accurately describe microglial heterogeneity in human health or disease [51]. To this end, integrative methodological approaches that can deeply phenotype microglia at the single-cell level have been employed to characterise microglial changes in ageing and disease. In the context of ALS, single cell and bulk RNA sequencing has been used to identify microglial states and assess their changes in rodent models and ALS patients [20,22,28–30,40,58–61], but their validation at the protein level in the human ALS brain has been limited. We recently utilised multiplexed immunohistochemistry to identify microglial changes relative to TDP-43 pathology in ALS [39]. We demonstrated that microglia have a phagocytic immunophenotype at early-stage ALS but transition to a dysfunctional immunophenotype at end-stage disease, and that these immunophenotypes are driven by TDP-43 aggregation [39], validating previous microglial states identified by RNA sequencing [40]. It is currently unclear how microglial morphologies change with respect to protein expression changes, but highly multiplexed immunohistochemistry enables the analysis of both concurrently, and the potential to infer changes in microglial function in the ALS brain.

This study aims to explore both microglial immunophenotype and morphology to characterise microglial states in ALS. We use automated single-cell protein expression and morphological analyses of immunohistochemically labelled tissue to compare changes in microglial immunophenotype and morphology in ALS brains and age-matched normal controls. We also infer temporal changes in microglial immunophenotype and morphology in ALS by examining TDP-43 mouse model brain tissue at disease onset, early disease, and late disease stages [62]. Our data demonstrate that microglia alter their morphologies parallel to pTDP-43 aggregation in ALS and that these alterations precede changes in levels of the microglial proteins HLA-DR, CD68, and Iba1. The immunophenotype and morphology data we present in this study support the hypothesis that microglial morphological alterations occur early in the ALS brain.

## Materials and methods

### Human case selection

Formalin-fixed paraffin-embedded motor cortex blocks from 10 neurologically normal and 10 ALS cases from the Neurological Foundation Human Brain Bank (HuBB) at the Centre for Brain Research, University of Auckland were used in this study (Table 1). Control cases had no previous history of neurological disorders and cause of death was unrelated to any neurological condition. ALS cases were diagnosed clinically by consultant neurologists at Auckland City or Middlemore Hospitals (Auckland, New Zealand) during life. All case classifications were confirmed with post-mortem neuropathology assessments performed by consultant neuropathologists at Auckland City Hospital. All ALS cases showed pTDP-43 deposition in the motor cortex. ALS cases were further classified as stage 1-3 or stage 4 pTDP-43 pathology based on the absence or presence of pTDP-43 aggregates in the hippocampus, respectively [63].

**Table 1.** Post-mortem case demographics.

| Table 1 Post-mortem case demographics |  |  |  |  |  |  |
| --- | --- | --- | --- | --- | --- | --- |
| Group |  | Case code | Age (y) | Sex | PMD (h) | fALS/sALS |
| Neurologically Normal Controls<br>(n = 10) |  | H211 | 41 | M | 8 | - |
|  |  | H215 | 67 | F | 23.5 | - |
|  |  | H226 | 73 | F | 48 | - |
|  |  | H230 | 57 | F | 32 | - |
|  |  | H238 | 63 | F | 16 | - |
|  |  | H240 | 73 | M | 26.5 | - |
|  |  | H242 | 61 | M | 19.5 | - |
|  |  | H245 | 63 | M | 20 | - |
|  |  | H247 | 51 | M | 31 | - |
|  |  | H249 | 77 | F | 17 | - |
| Control mean ± SD |  | - | 62.60 ± 10.95 | 5M:5F | 24.15 ± 11.06 | - |
| ALS<br>(n = 10) | Stage 1-3<br>(n = 5) | MN11 | 77 | F | 18 | fALS<br>(unknown) |
|  |  | MN14 | 59 | F | 19 | fALS<br>(unknown) |
|  |  | MN16 | 69 | M | 16.5 | sALS |
|  |  | MN20 | 85 | M | 15 | sALS |
|  |  | MN25 | 71 | M | 23.5 | sALS |
|  | Stage 4<br>(n = 5) | MN13 | 55 | M | 10 | sALS<br>(ANXA11 p.G38R) |
|  |  | MN15 | 54 | F | 18 | sALS |
|  |  | MN19 | 75 | M | 20.5 | sALS |
|  |  | MN29 | 72 | M | 24 | sALS |
|  |  | MN30 | 84 | F | 17.5 | sALS |
| ALS mean ± SD |  | - | 70.10 ± 11.07 | 6M:4F | 18.20 ± 4.063 | - |

The fixation and dissection of anatomical regions from human brains by the HuBB has been described previously [64]. Briefly, human brains were obtained at autopsy and the right hemisphere was fixed by perfusion of 15% formaldehyde in 0.1 M phosphate buffer through the cerebral arteries. Brains were subsequently dissected into approximately 60 regional blocks and 1-cm thick blocks from each region were processed for paraffin embedding. For this study, 10-µm thick sections were cut from paraffin-embedded motor cortex blocks from all control and ALS cases (Table 1) using a rotary microtome and mounted on ÜberFrost® Printer Slides (InstrumeC).

### Multiplexed fluorescence immunohistochemistry and imaging of human ALS brain

A 5-plex immunohistochemistry labelling panel was designed to quantify microglial immunophenotypes (anti-CD68, -HLA-DR, and -Iba1) and morphology (anti-Iba1) relative to pathology (pTDP-43) per cell (Hoechst nuclear stain) (Table 2, quantification panel). CD68 and Iba1 were subsequently co-labelled to validate their expression by microglia with specific morphologies (Table 2, confocal panel).

**Table 2.** Immunohistochemistry panels for analysis of microglia changes in the human ALS brain.

|  |  |  |  |
| --- | --- | --- | --- |
| **PANEL 1** Quantification panel |  |  |  |
| **Primary antibody,**<br>**Concentration** | **Company,**<br>**Catalogue**<br>**number** | **Secondary antibody,**<br>**Concentration** | **Company, Catalogue**<br>**number** |
| Rabbit IgG anti-pTDP-43<br>(Ser 409/410), 1:3,000 | Cosmobio, TIP-PTD-P02 | Goat anti-rabbit<br>AlexaFluor® 488, 1:500 | Jackson<br>Laboratories, 111-545-144 |
| Mouse IgG<sub>1</sub> anti-HLA-DR,<br>1:1,000 | DAKO, M0775 | Goat anti-mouse IgG1<br>AlexaFluor® 546, 1:500 | Thermo Fisher<br>Scientific, A21123 |
| Mouse IgG<sub>3</sub> anti-CD68,<br>1:200 | Abcam, ab783 | Goat anti-mouse IgG3<br>AlexaFluor® 594, 1:500 | Thermo Fisher<br>Scientific, A21155 |
| Guinea pig IgG anti-Iba1,<br>1:2,000 | Synaptic<br>Systems, 234-004 | Goat anti-chicken<br>AlexaFluor® 647, 1:500 | Thermo Fisher<br>Scientific, A21450 |
| **PANEL 2** Confocal panel |  |  |  |
| **Primary antibody,**<br>**Concentration** | **Company,**<br>**Catalogue**<br>**number** | **Secondary antibody,**<br>**Concentration** | **Company, Catalogue**<br>**number** |
| Chicken IgY anti-Iba1,<br>1:1,000 | Synaptic<br>Systems, 234-009 | Goat anti-chicken<br>AlexaFluor® 488, 1:500 | Thermo Fisher<br>Scientific, A11039 |
| Rabbit IgG anti-CD68,<br>1:1,000 | Abcam,<br>ab215212 | Goat anti-rabbit<br>AlexaFluor® 594, 1:500 | Jackson<br>Laboratories, 115-585-166 |

Immunohistochemical labelling of paraffin-embedded motor cortex sections was carried out as previously described [39,65,66]. Briefly, Tris-EDTA pH 9.0 antigen retrieval was performed, and sections were incubated in primary then secondary antibody mixtures (Table 2). Sections were washed thoroughly in phosphate-buffered saline between each step. Nuclei were counterstained with Hoechst 33258 and sections were coverslipped using ProLong® Gold Antifade mounting media.

Sections labelled with the 5-plex immunohistochemical panel were imaged on a Zeiss Z2 Axioimager (20×/0.9 NA) using MetaSystems VSlide acquisition software and MetaCyte stitching software. The Zeiss Z2 Axioimager used a Colibri 7 solid-state light source with LED lamps and the following filter sets to enable spectral separation of the 5 fluorophores per round (Ex peak (nm); Em peak (nm)/bandpass (nm)): Hoechst 33258 (385; 447/60), AlexaFluor® 488 (475; 550/32), AlexaFluor® 546 (555; 580/23), AlexaFluor® 594 (590; 628/32), and AlexaFluor® 647 (630; 676/29).

Sections labelled with anti-CD68 and -Iba1 only were imaged on a Zeiss LSM 800 Airyscan confocal microscope (63x/1.4 NA, oil immersion) using ZEN 2.6 software (Zeiss). Optical *z*-stacks were taken through the entirety of the cell body and processes. Images were acquired using the built-in Airyscan module and processed using the ZEN microscopy software (Zeiss). Maximum intensity *z*-projections were generated and processed using FIJI (v 1.53C).

### Multiplexed fluorescence immunohistochemistry and imaging of transgenic mouse model ALS brain

Brain tissue from a bigenic *NEFH*-tTA/*tetO*-hTDP-43ΔNLS (rNLS) mouse model of ALS (and litter-matched controls) at disease onset, early disease, and late disease stages (2, 4, and 6 weeks off doxycycline (WOD), respectively) was immunohistochemically labelled with CD68, Iba1, NeuN, and pTDP-43 and imaged on a Zeiss Z2 Axioimager (20×/0.9 NA) using MetaSystems VSlide acquisition software and MetaCyte stitching software (as described above) [39]. The images from this cohort of rNLS mice have been analysed as part of a previous study [39], but have been reanalysed here and new data presented.

### Quantification of microglial features and ALS pathology using MetaMorph image analysis pipelines

To quantify microglial protein expression levels and morphological features in the human and transgenic mouse model ALS brain, custom image analysis pipelines were developed in Metamorph software (Molecular Devices), similar to those previously described [39,67,68]. Prior to all analyses, manual regions of interest (ROI) were drawn on each Hoechst image to isolate cortical layers I-VI for analysis and exclude tissue folds and defects.

#### Tissue-wide microglial and pTDP-43 integrated intensity analyses

To identify global microglial and pTDP-43 changes in the ALS motor cortex, the tissue-wide abundance of each microglial marker and pTDP-43 was measured. For the human tissue, binary masks of CD68, HLA-DR, and Iba1 immunolabelling were generated using the adaptive thresholding tool. These three binary masks were combined to create a master mask, encompassing all microglia immunoreactive for CD68, HLA-DR, and/or Iba1. For the mouse tissue, a binary mask of only Iba1 was generated to identify all microglia as all microglia were highly immunoreactive for Iba1. The integrated intensity of each marker was measured across the master mask (human) or Iba1 mask (mouse) and normalised to the ROI area to give a measure of tissue-wide expression. To measure the tissue-wide abundance of pTDP-43, a binary mask of specific pTDP-43 immunoreactivity was generated using the threshold clip tool. The integrated intensity of pTDP-43 labelling within this binary mask was measured and normalised to ROI area.

#### Tissue-wide microglial morphology analyses

To identify overall microglial morphology changes in the ALS motor cortex, the tissue-wide averages of key cell morphology features were measured: cell body area, process number, total outgrowth, and branch number. These cell morphology features were measured from the Iba1 image using the Metamorph in-built Neurite Outgrowth application. This application identified microglial cell bodies based on manually determined maximum cell body width, cell body Iba1 intensity above local background, and minimum cell body area, and microglial processes based on maximum process width, process Iba1 intensity above local background, and minimum process length to be considered significant.

#### Single-cell microglial immunophenotype and morphology analyses

Next, all microglia within each ROI were identified and the single-cell expression of CD68, HLA-DR, and Iba1 and cell morphology was measured. Using the microglial master mask generated for the tissue-wide microglial integrated intensity analysis, the watershed lines morphology filter was used to identify the border of each microglial domain; each microglia and its domain was contained within its own border. To quantify the single-cell expression of each microglial marker, the average intensity of CD68, HLA-DR, and Iba1 within the master mask was measured in each microglial domain. To quantify the single-cell morphology features, cell body area, process number, total outgrowth, and branch number were measured from Iba1 using the Neurite Outgrowth application in each microglial domain.

### Microglial phenotype and morphological feature clustering

To identify potential clusters of microglial immunophenotypes or morphologies, cells were evenly sampled across the different groups. For the human data, 7,340 cells each were randomly sub-sampled from stage 1-3 ALS, stage 4 ALS, and normal cases for a total of 22,020 cells. The single-cell morphological features of mean microglial cell body area, process number, total outgrowth, and branch number in addition to mean cell intensities of Iba1 and CD68 were included in our dataset. These were log-transformed for equal weighting and a k-nearest neighbor (KNN) graph was constructed with the k parameter set as 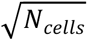. For the cell clustering, a community detection clustering approach (Louvain clustering) was applied with the resolution (clustering granularity) set to 1.0 wherein a total of 11 cluster assignments were obtained. All data processing and subsequent clustering was conducted in R v.4.2 using the ‘FNN v.1.1.3’ and ‘igraph v.1.2.11’ packages [69,70].

### Statistical analyses

For human analyses, pTDP-43 and microglial marker tissue-wide integrated intensities and microglial morphology measures were compared between case groups with multiple Mann-Whitney tests and multiple comparisons were controlled for using a False Discovery Rate of 0.01, as determined by the two-stage step-up method of Benjamini, Krieger, and Yekutieli. Statistical significance was set at p ≤ 0.05 with significances of difference between case groups shown as *p ≤ 0.05, **p ≤ 0.01, ***p ≤ 0.001, ****p≤ 0.0001. PTDP-43 and microglial marker tissue-wide integrated intensities and microglial morphology measures were sequentially correlated with one another using Spearman correlations. When r ≤ −0.7 or r ≥ 0.7 and p ≤ 0.05, correlations were considered statistically significant and strong. When −0.7 < r ≤ −0.4 or 0.7 > r ≥ 0.4 and p ≤ 0.05, correlations were considered statistically significant and moderate. For the microglial clustering analysis, the percentage of microglia in each cluster was compared between case groups using a 2-way ANOVA with Tukey’s multiple comparisons test, and the mean single cell CD68 and Iba1 average intensities were compared between clusters using a Kruskal-Wallis test with Dunn’s multiple comparisons test.

For mouse analyses, microglial intensity and morphology measures were compared between case groups at each time point with 2-way ANOVA with Tukey’s multiple comparisons test. Statistical significance was set at p ≤ 0.05 with the significance of differences between groups shown as *p ≤ 0.05, **p ≤ 0.01, ***p ≤ 0.001, ****p≤ 0.0001.

## Results

### Increased microglial CD68 and Iba1 levels and pTDP-43 abundance in the human ALS motor cortex

Immunohistochemistry was used to identify microglial markers HLA-DR, CD68, and Iba1, and pTDP-43 aggregates, in the control and ALS motor cortex including cases previously shown to have pTDP-43 in defined regions including the hippocampus (representing more advanced disease, ‘stage 4 ALS’) [63] and cases with pTDP-43 in the motor cortex and/or spinal cord but not hippocampus (representing less advanced disease, ‘stage 1-3 ALS’, Fig 1A-E). Tissue-wide integrated intensity analyses revealed that microglial HLA-DR intensity remained unchanged in ALS (Fig 1F), while CD68 and Iba1 intensities were significantly increased in both stage 1-3 ALS and stage 4 ALS compared with controls (Fig 1G and H). As expected, we observed no pTDP-43 pathology in the motor cortex of control cases (Fig 1B and I), with pTDP-43 only seen in ALS cases (Fig 1C, E, and I). However, a statistically significant increase in pTDP-43 compared with controls was only seen in stage 4 ALS cases, likely due to the variability in pTDP-43 pathology levels in stage 1-3 cases (Fig 1I).

**Figure 1:**
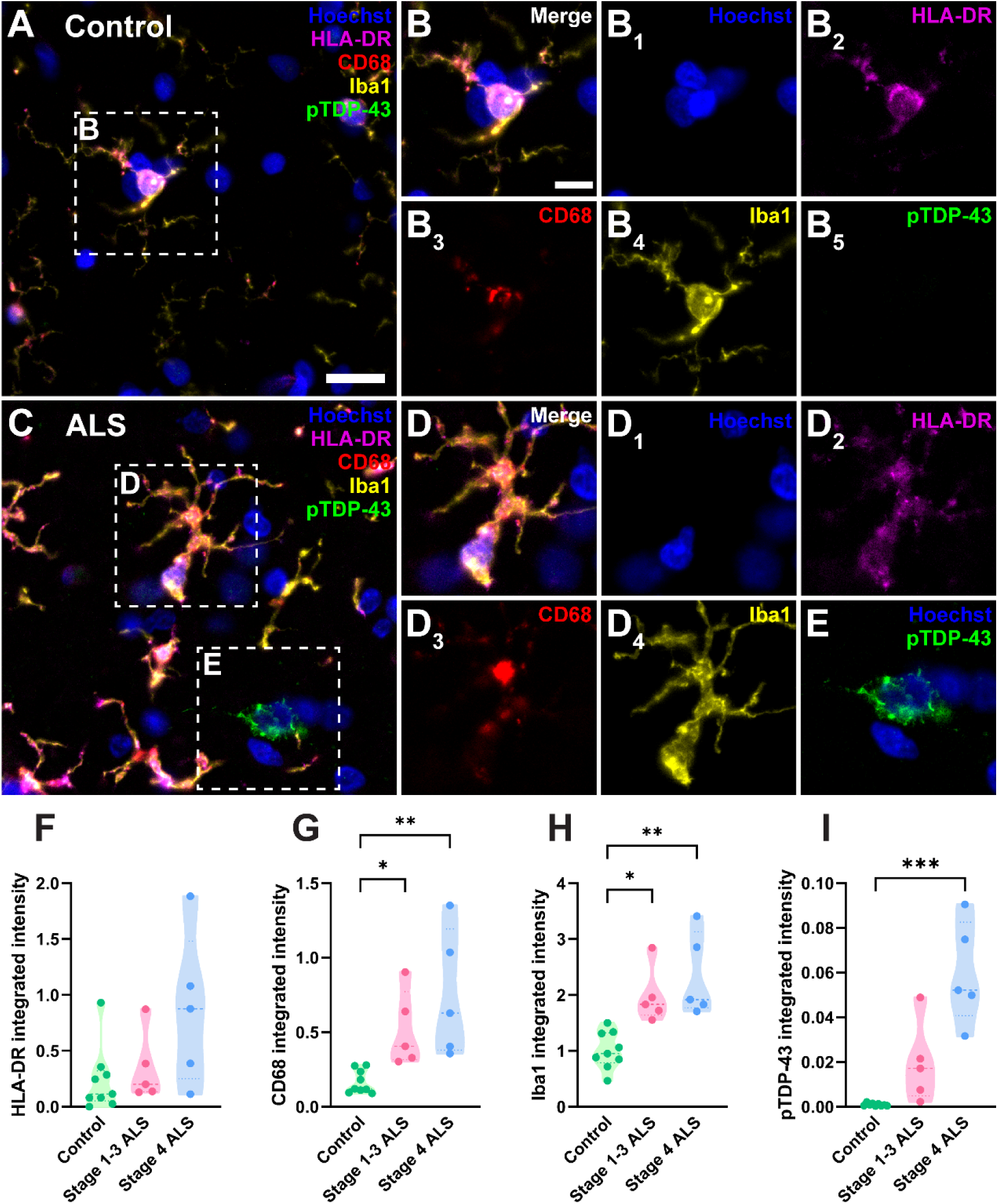
Increased microglial CD68 and Iba1 levels and pTDP-43 abundance in the human ALS motor cortex. Immunohistochemical labelling was used to visualise microglia and pTDP-43 in the motor cortex from human control (A-B), stage 1-3 (not shown), and stage 4 ALS (C-E) cases. Microglial markers, HLA-DR (magenta), CD68 (red), and Iba1 (yellow), were co-labelled with pTDP-43 (green) and a Hoechst nuclear counterstain (blue). Scale bar (A and C) = 20 µm and scale bar (B, D, and E) = 10 µm. Total microglia were identified by creating separate binary masks from thresholded HLA-DR, CD68, and Iba1 images which were then combined to create a microglial master mask [39,68,80]. The total integrated intensities of HLA-DR (F), CD68 (G), and Iba1 (H) were measured within this master mask, normalised to tissue area and compared between control, stage 1-3 ALS, and stage 4 ALS cases. The total integrated intensity of pTDP-43 aggregates normalised to tissue area was quantified and compared between normal, stage 1-3 ALS, and stage 4 ALS cases (I). Data presented as truncated violin plots with median and quartiles shown; control n = 10, stage 1-3 ALS n = 5, and stage 4 ALS n = 5. All intensity measures were compared between case groups with multiple Mann-Whitney tests and multiple comparisons were controlled for using a False Discovery Rate of 0.01, as determined by the two-stage step-up method of Benjamini, Krieger, and Yekutieli. Significances of difference between case groups: *p ≤ 0.05, **p ≤ 0.01, ***p ≤ 0.001.

### Microglial morphologies change in human ALS and correlate with CD68 and Iba1 levels

We subsequently sought to identify whether microglial morphologies were altered in the human ALS motor cortex. Microglial cell bodies and processes were identified by Iba1 immunoreactivity using the Metamorph Neurite Outgrowth application (Fig 2A), as previously described [71]. Single-cell morphology measurements of cell body area, process number, total outgrowth, and branch number were averaged across all microglial cells in each case to give a mean measure per microglia (Fig 2B-E). We identified significant changes in microglial morphology measures in ALS: cell body area and process number were increased in stage 1-3 ALS compared with controls (Fig 2B and C), and total outgrowth was significantly reduced in stage 4 ALS compared with controls (Fig 2D).

**Figure 2:**
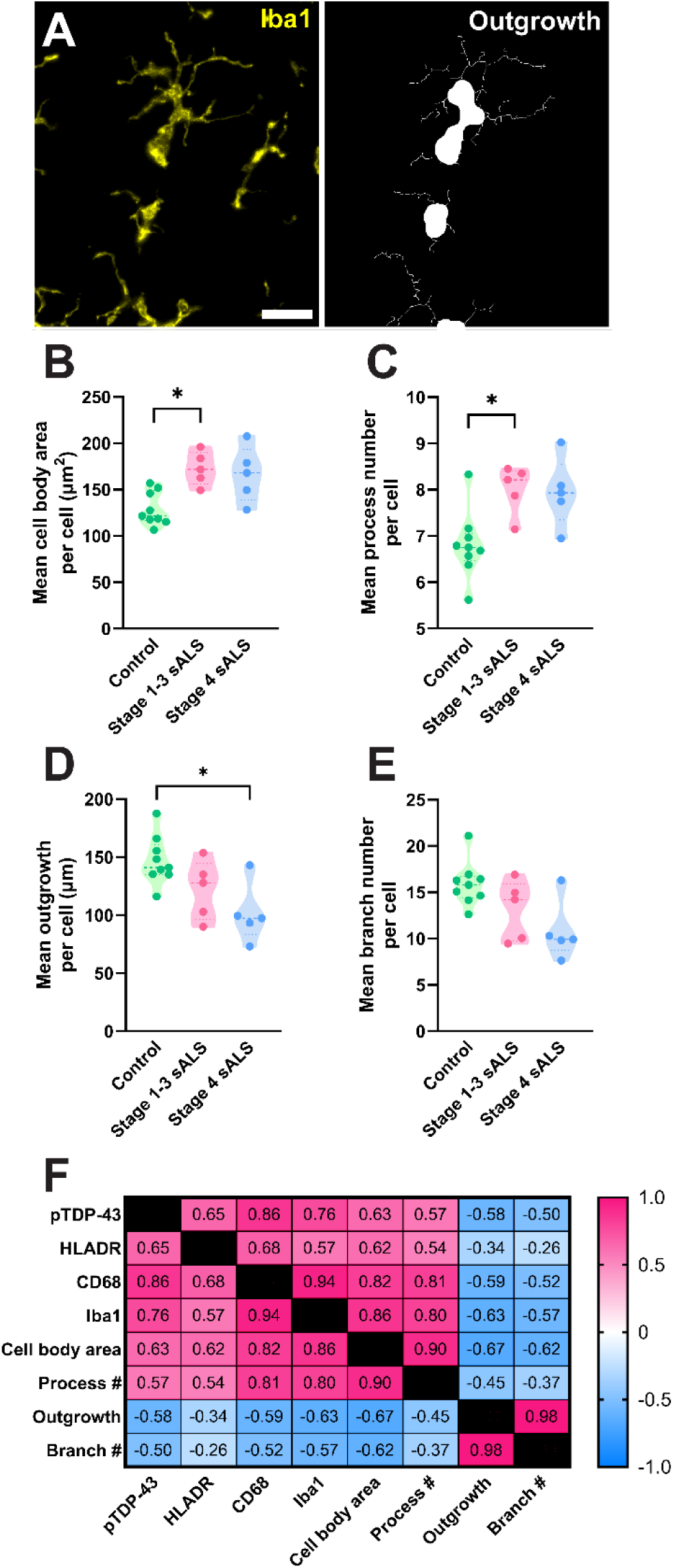
Microglial morphologies change in human ALS and correlate with CD68 and Iba1 levels. Microglial cell bodies and processes were identified using Iba1 immunoreactivity (A, yellow) and used to quantify morphology measures per cell (A, white): cell body area (B), process number (C), outgrowth (D), and branch number (E). Single-cell measures were averaged across all microglia in each case to give a mean measure per cell which was compared between control, stage 1-3 ALS, and stage 4 ALS cases. Data presented as truncated violin plots with median and quartiles shown; control n = 10, stage 1-3 ALS n = 5, and stage 4 ALS n = 5. All morphology measures were compared between case groups with multiple Mann-Whitney tests and multiple comparisons were controlled for using a False Discovery Rate of 0.01, as determined by the two-stage step-up method of Benjamini, Krieger, and Yekutieli. Significances of difference between case groups: *p ≤ 0.05. Measures of pTDP-43 pathology, microglial marker intensities, and microglial morphology were sequentially correlated in all cases (n = 20) using Spearman correlations (F). The resulting r value from each correlation is presented in the correlation matrix and colour coded relative to strength. When r ≤ −0.7 or r ≥ 0.7 and p ≤ 0.05, correlations were considered statistically significant and strong. When −0.7 < r ≤ −0.4 or 0.7 > r ≥ 0.4 and p ≤ 0.05, correlations were considered statistically significant and moderate.

To identify whether microglial morphological changes correlated with pTDP-43 pathology and/or microglial HLA-DR, CD68, and Iba1 levels, the mean morphology measures above were sequentially correlated with tissue-wide pTDP-43, HLA-DR, CD68, and Iba1, integrated intensities (Fig 2F and S1). In accordance with the pattern of changes in the human ALS motor cortex (Fig 1), pTDP-43, CD68, and Iba1 showed significant strong positive correlations with one another (Fig 2F). PTDP-43, CD68, and Iba1 also all showed significant correlations with all morphology measures: moderate to strong positive correlations with cell body area and process number, and moderate negative correlations with total outgrowth and branch number. Furthermore, morphological measures significantly correlated with one another (Fig 2F and S1). This suggests that as microglial CD68 and Iba1 levels increase in ALS with increased pTDP-43 aggregation, microglia also reduce their outgrowth and branch number, and increase their process number and cell body area, suggestive of hypertrophic morphology.

### Microglial clusters enriched in human ALS are characterised by high Iba1 and CD68 levels with hypertrophic and dystrophic morphologies

Having determined that microglial morphology and functional marker expression are altered in ALS and are correlated, we sought to identify specific microglial phenotypes emerging in ALS. To identify microglial phenotypes enriched in ALS cases, microglia from control, stage 1-3 ALS, and stage 4 ALS cases were clustered based on the single cell CD68, HLA-DR, and Iba1 average intensities, and morphological measures (cell body area, process number, total outgrowth, and branch number). A total of 11 unique clusters were identified, with 4 clusters (numbered arbitrarily) being differentially abundant in ALS compared with control cases (Fig 3A). Clusters 1 and 8 were depleted in ALS cases: cluster 1 was significantly more abundant in control cases compared with stage 4 ALS cases, and cluster 8 was significantly more abundant in control cases compared with both stage 1-3 ALS and stage 4 ALS cases (Fig 3A). However, no significant correlations between the abundance of control- or ALS-enriched clusters and pTDP-43 intensity were identified (Fig 3B and C).

**Figure 3:**
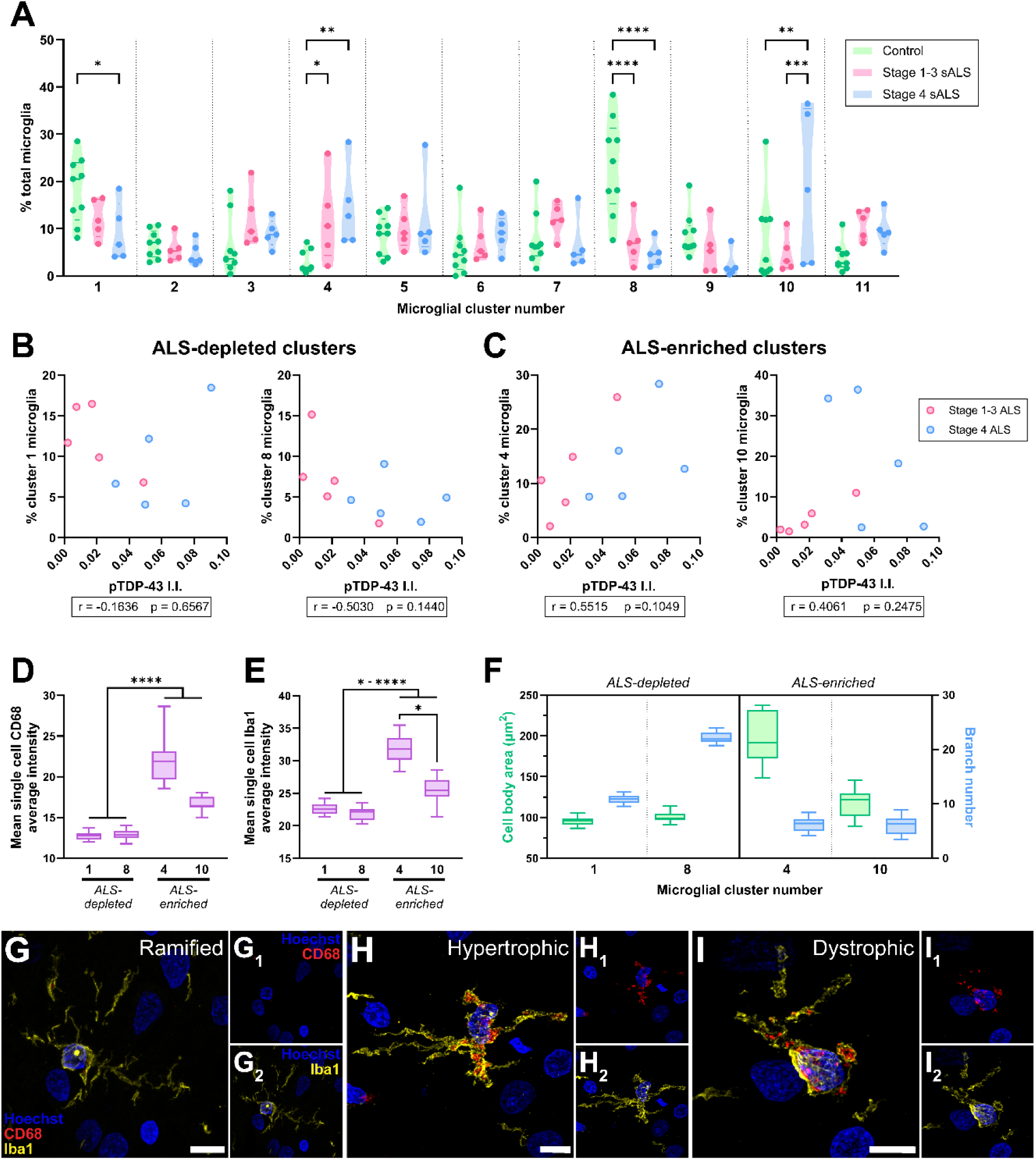
Microglial clusters enriched in human ALS are characterised by high cell body area, low branch number, and high Iba1 and CD68 levels. Louvain clustering was carried out using the single-cell average intensities of CD68, HLA-DR, and Iba1, and morphological measures (cell body area, process number, outgrowth, and branch number) from a randomly subsampled 7,340 microglia per case group (22,020 cells total), resulting in 11 unique clusters (arbitrarily numbered) (**A**). The percentage of microglia in each cluster was compared between control, stage 1-3 ALS, and stage 4 ALS cases using a 2-way ANOVA with Tukey’s multiple comparisons test; data presented as truncated violin plots with median and quartiles shown; control n = 10, stage 1-3 ALS n = 5, and stage 4 ALS n = 5. For clusters depleted (1 and 8; **B**) and enriched (4 and 10; **C**) in ALS cases, the percentage of microglia in each cluster was correlated with pTDP-43 integrated intensity (I.I.) using Spearman’s correlation; each correlation was carried out using a total n = 10 cases, with stage 1-3 ALS and stage 4 ALS cases colour-coded and r and p values presented for each correlation. The mean single-cell CD68 and Iba1 average intensity was compared between clusters using a Kruskal-Wallis test with Dunn’s multiple comparisons test (**D** and **E**); data presented for ALS-depleted and ALS-enriched clusters as a box and whisker graph, with minimum, maximum, and median shown (n = 20). The mean cell body area and single-cell branch number were determined for all clusters (**F**); data presented for ALS-depleted and ALS-enriched clusters as a box and whisker graph, with minimum, maximum, and median shown (n = 20). Significance of differences between groups: *p ≤ 0.05, **p ≤ 0.01, ***p ≤ 0.001, ****p ≤ 0.0001. Confocal microscopy was used to visualise ramified (G), hypertrophic (H), and dystrophic microglial morphologies (I); example images from stage 4 ALS case, MN29, are shown; scale bars = 10 µm.

We subsequently phenotyped the ALS-depleted and ALS-enriched microglial clusters, characterising their CD68 and Iba1 levels and morphological characteristics (Fig 3D-F). The ALS-enriched clusters 4 and 10 had significantly higher CD68 and Iba1 average intensities than the ALS-depleted clusters (Fig 3D and E) or the clusters equally present in ALS and control cases (Fig S2). Furthermore, ALS-enriched cluster 4 had significantly higher CD68 and Iba1 average intensities than ALS-enriched cluster 10 (Fig 3D and E). ALS-depleted and ALS-enriched clusters also showed unique morphological characteristics. Only cell body area and branch number and were assessed because the mean cell body area showed a strong significant correlation with process number (r = 0.85, p < 0.0001), and branch number in each cluster showed a strong significant correlation with total outgrowth (r = 0.99, p < 0.0001). ALS-depleted clusters 1 and 8 had smaller cell bodies with high branch numbers, reminiscent of a ramified morphology (Fig 3F and S3), implying depletion of ramified microglia in ALS. Conversely, ALS-enriched clusters had larger cell bodies with low branch numbers with reminiscent of hypertrophic and dystrophic morphologies (Fig 3F and S3). We confirmed the presence of CD68^low^ Iba1^low^ ramified microglia (clusters 1 and 8) and CD68^high^ Iba1^high^ hypertrophic and dystrophic microglia (clusters 4 and 10) in the control and ALS motor cortex using confocal microscopy (stage 4 ALS case example images shown in Fig 3G-I).

### Microglial Iba1 and morphological changes occur parallel to pTDP-43 aggregation and prior to CD68 changes in the TDP-43-driven rNLS mouse model of ALS

With changes in microglial marker expression and morphological features, we sought to determine *when* these microglial changes occur with respect to the development of pTDP-43 aggregation. To understand this relationship temporally, we utilised brain tissue from bigenic *NEFH*-tTA/*tetO*-hTDP-43ΔNLS (rNLS) and single transgenic *tetO*-hTDP-43ΔNLS (control) mice at 2, 4, and 6 weeks off doxycycline (WOD), equivalent to disease onset, early disease, and late disease stages, respectively [62].

We first investigated the presence of ramified, hypertrophic, and dystrophic microglia in the rNLS mouse motor cortex. Indeed, as per the human ALS motor cortex, we identified all three microglial morphologies in the rNLS motor cortex at 6 WOD (Fig 4A-C). Furthermore, hypertrophic and dystrophic microglia showed increased CD68 expression (Fig 4B and C), as per human microglia clusters 4 and 10.

**Figure 4:**
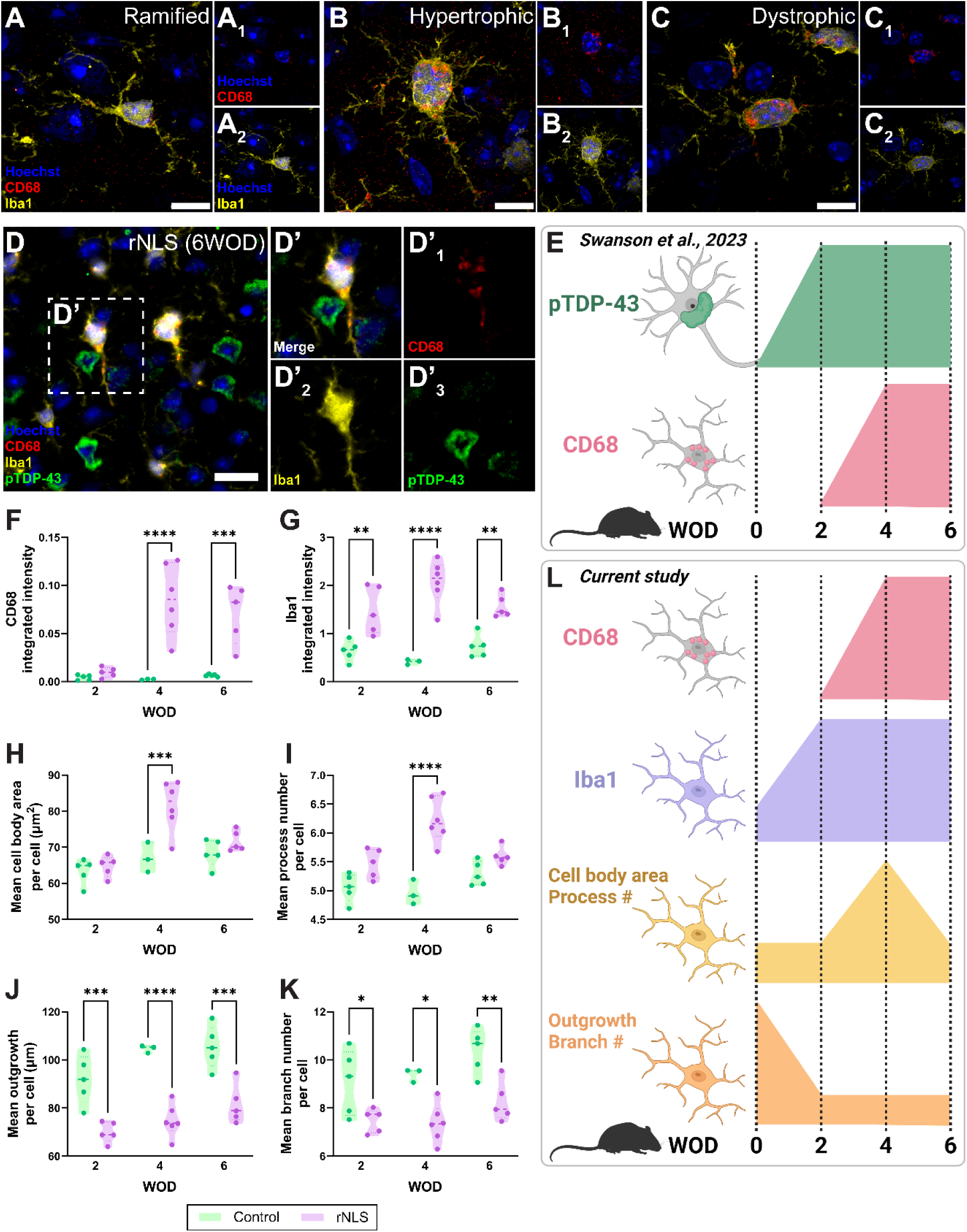
Microglial Iba1 and morphological changes occur parallel to pTDP-43 aggregation and prior to CD68 changes in the TDP-43-driven rNLS mouse model of ALS. Immunohistochemical labelling of Iba1 and CD68 was carried out on motor cortex tissue from bigenic NEFH-tTA/tetO-hTDP-43ΔNLS (rNLS) and single transgenic tetO-hTDP-43ΔNLS (control) mice at 2, 4, and 6 weeks off DOX (WOD). Confocal microscopy was used to visualise ramified (A), hypertrophic (B), and dystrophic microglial morphologies; example images from rNLS at 6 WOD are shown; Scale bars = 10 µm. Images previously analysed in Swanson et al., (2023) were reanalysed to investigate temporal changes in microglial morphology measures relative to pTDP-43 aggregation; an example immunohistochemical image of CD68 (red), and Iba1 (yellow), pTDP-43 (green) and a Hoechst nuclear counterstain (blue) labelling from a rNLS mouse at 6 WOD is shown (D); scalebar = 20 µm. Summary of Swanson et al., (2023) findings is shown in E; created with BioRender.com. Total microglia were identified by creating separate binary masks from CD68 and Iba1 images which were then combined to create a microglial master mask. The integrated intensities of CD68 (F) and Iba1 (G) were measured within this master mask and compared between control and rNLS mice at 2, 4, and 6 WOD. Microglial cell bodies and processes were identified using Iba1 immunoreactivity and used to quantify morphology measures per cell: cell body area (H), process number (I), outgrowth (J), and branch number (K). Single-cell measures were averaged across all microglia in each case to give a mean measure per cell which was compared between control and rNLS mice at 2, 4, and 6 WOD. Data presented as truncated violin plots with median and quartiles shown; control n = 3-5, rNLS n = 4-6. Each intensity and morphology measure was compared between case groups at each WOD timepoint with 2-way ANOVA with Tukey’s multiple comparisons test. Significance of differences between case groups: *p ≤ 0.05, **p ≤ 0.01, ***p ≤ 0.001, ****p ≤ 0.0001. Summary of current study’s findings is shown in L; created with BioRender.com.

To quantify microglial morphological features relative to CD68 and Iba1 levels and neuronal pTDP-43 levels, we utilised images from a previous study [39] (Fig 4D). Using these images, we have previously shown an increase in the percentage of CD68^high^ microglia at 4 WOD, occurring after the accumulation of pTDP-43 from 2 WOD (Fig 4E). In the current study, we identified a significant increase in CD68 integrated intensity at 4 and 6 WOD in rNLS mice relative to controls (Fig 4F). In addition, Iba1 integrated intensity was significantly increased from 2 WOD in the rNLS mouse cohort relative to controls (Fig 4G), suggesting microglial changes occur prior to the CD68 changes observed at 4 WOD.

Changes in microglial morphological measures were also identified from 2 WOD in the rNLS mice (Fig 4H-K). At 2 WOD, rNLS mouse microglia had unchanged cell body area and process number, but significantly reduced total outgrowth and branch number, suggesting a change towards a hypertrophic morphology (Fig 4H-K). At 4 WOD, rNLS mouse microglia had significantly increased cell body area and process number and significantly reduced total outgrowth and branch number, suggesting predominantly hypertrophic morphologies (Fig 4H-K). Finally, at 6 WOD, rNLS mouse microglia resembled 2 WOD microglia, with unchanged cell body area and process number, and significantly reduced total outgrowth and branch number, suggesting microglia become less hypertrophic and more dystrophic (Fig 4H-K). These data are summarised in Fig 4L.

Together these data suggest microglial changes occurred in parallel with neuronal pTDP-43 accumulation; at disease-onset, microglia increased Iba1 expression and changed towards a hypertrophic morphology, at early-disease stage, microglia increased CD68 expression and became hypertrophic, and at late-stage disease, microglia became dystrophic.

## Discussion

In this study, we used fluorescent immunohistochemistry and single cell analysis pipelines to identify changes in microglial immunophenotypes and morphologies relative to pTDP-43 aggregation in the ALS motor cortex. Microglial subtypes enriched in human ALS were defined by high CD68 and Iba1 levels and hypertrophic and dystrophic morphologies. Indeed, similar microglial changes were identified in the rNLS mouse model of ALS following the onset of pTDP-43 aggregation. Overall, this study provides an examination of microglial immunophenotype and morphological changes in human ALS following pTDP-43 aggregation.

The increases in microglial CD68 and Iba1 in the ALS motor cortex significantly correlated with neuronal pTDP-43 pathology load (Fig 1). The identified increase in CD68 corroborates our previous work where we identified a robust increase in CD68^high^ microglia in the human ALS motor cortex that correlated with pTDP-43 pathology load [39]. Because CD68 is classically considered a marker of phagocytic microglia, in the context of ALS this increase in microglial CD68 expression is hypothesised to be a reaction to the phagocytosis of pTDP-43 aggregates [20,72–75]. The tissue-wide increase in Iba1 identified in this study also corroborates our previous work [39]. Iba1 is a cytoskeletal-associated calcium binding protein involved in membrane ruffling and phagocytosis [76–79]. Previous studies have described increases in Iba1 expression by hypertrophic, reactive microglia in the normal and *SOD1* ALS brain [15,17,18,21]. Indeed, both CD68 and Iba1 expression correlated with tissue-wide microglial morphology measures suggesting both markers are up-regulated by hypertrophic, reactive microglia (Fig 2).

We subsequently carried out single cell analyses to better elucidate the relationship between CD68 and Iba1 expression levels and microglial morphologies. We identified CD68^high^ Iba1^high^ microglial subtypes with hypertrophic and dystrophic morphologies enriched in the human ALS motor cortex (Fig 3). Hypertrophic microglia are considered classically reactive, with an enlarged cell body to increase metabolic activity and shortened processes to explore their immediate surroundings [50]. Dystrophic microglia show evidence of cytoplasmic fragmentation, similar to that seen in apoptotic cells, and are considered to be diseased or dysfunctional microglia [44,50]. We therefore postulate that, in the human ALS brain, microglia change their morphologies at different points in response to pTDP-43: first, microglia increase Iba1 expression and become hypertrophic to increase metabolic activity and phagocytose pTDP-43, resulting in increased CD68 expression. Later, microglia become ‘exhausted’ from chronic reactions to ALS-associated pathology and degeneration, resulting in dysfunction and a dystrophic morphology. Our previous study identified high expression of the dysfunction marker, L-ferritin, in ALS-enriched clusters [39] and the specific expression of L-ferritin by dystrophic microglia in the human brain [42] validates the presence of microglial dysfunction in ALS.

Arguing against pTDP-43 pathology driving microglial immunophenotype and morphology changes in the ALS brain was the lack of significant correlations between the abundances of CD68^high^ Iba1^high^ hypertrophic and dystrophic microglial subtypes and pTDP-43 load in this study (Fig 3). This could be because large end-stage TDP-43 inclusions are unlikely to be the ‘toxic species’ in ALS, and the analysis pipeline used to measure pTDP-43 pathology load will not detect the other aggregation pathway species that *are* likely toxic; misfolded monomers and oligomers [38]. Comparison of features between stage 1-3 and stage 4 ALS cases classified on pTDP-43 load provides some insight to compare disease at different stages of progression. However, one of the primary limitations of assessing cellular changes in post-mortem human tissue is the inability to assess temporal changes with certainty. To overcome this limitation, we utilised brain tissue from the rNLS TDP-43 mouse model of ALS (*NEFH*-tTA/*tetO*-hTDP-43^ΔNLS^); the removal of dietary DOX induces the cytoplasmic accumulation of TDP-43 in neurons, causing motor neuron death in the spinal cord and motor cortex and associated motor phenotypes [62]. We assessed microglial immunophenotype and morphology changes at 2, 4, and 6 WOD, equivalent to disease onset, early disease, and late disease stages, respectively (Fig 4). In our previous study, we identified a significant increase in pTDP-43 aggregate load by 2 WOD, with microglial CD68 levels increasing at 4 WOD [39]. In this study, we identified that microglia increase Iba1 expression and change to a hypertrophic morphology by 2 WOD (in parallel with pTDP-43 aggregation), microglia then increase CD68 expression at 4 WOD, and change to a dystrophic morphology at 6 WOD. These data support our hypotheses based on human tissue findings: that microglia change morphologies to aid in their phagocytosis of pTDP-43 but chronic reactivity results in dysfunction and dystrophy.

This study used a refined immunofluorescent panel of three commonly used microglial functional markers (HLA-DR, CD68, and Iba1) with single-cell analysis pipelines to assess microglial immunophenotype and morphology changes in ALS. However, neither these markers nor morphologies fully capture microglial heterogeneity of the human brain in health or disease. Iba1 and HLA-DR were not expressed in all microglial states in the normal and Alzheimer’s disease brain [42,80–82]. In the context of ALS, our previous study identified a tissue-wide increase in Iba1 levels, but Louvain clustering of single cell immunophenotypes did not reveal a higher expression of Iba1 in microglial states enriched in the ALS motor cortex [39]. However, in the current study where microglial morphologies were quantified and included as metrics in single-cell clustering, Iba1 was expressed more highly in ALS-enriched microglial states; likely a reflection of the functional link between Iba1 and morphology changes. Therefore, the more aspects of microglial phenotypes accounted for by analysis, the greater the depth of data, and the more specific disease-associated states identified. We have demonstrated that multiplexed immunohistochemistry and image analysis approaches can be integrated to identify microglial heterogeneity and subtle microglial state changes in disease.

The primary limitation of this study is that we are predicting changes in microglial function in ALS based on immunophenotype and morphology changes in post-mortem tissues. However, the analysis of microglial immunophenotypes and morphology in end-stage ALS patient tissue and a mouse model of ALS allows us to create a strong hypothesis of microglial functional changes in the ALS brain. However, one of the dominating narratives on microglia in the ALS field is the idea of microglia being neuroprotective in early-stage disease and neurotoxic in late-stage disease. This hypothesis was first proposed as microglia isolated from early-stage G93A *SOD1* mice were neurotrophic when co-cultured with neurons, whereas microglia from late-stage G93A *SOD1* mice were neurotoxic [31]. As a neurotoxic microglial phenotype can be induced by native and mutant forms of TDP-43 in vitro [32], it is hypothesized that this neurotrophic-to-neurotoxic change also happens in the 97% of ALS cases with TDP-43 pathology. Indeed, a recent study by Xie and colleagues demonstrated a neuroprotective microglial phenotype in rNLS mice and sporadic human ALS brains [41]; rod microglia emerge in at early-stage disease and have a neuroprotective transcriptomic signature. However, as Xie and colleagues do not describe the presence of hypertrophic and dystrophic microglial morphologies characteristic of microglia in human diseased brains, it is difficult to draw similarities to their study and our current study. The data presented in this study cannot confirm nor negate the neurotrophic-to-neurotoxic hypothesis; further functional studies investigating the effects of these immunophenotype and morphological changes are needed. However, we validate the presence of dysfunctional microglia in the end-stage disease ALS brain reflecting a potential loss of trophic support from microglia in late-stage disease. Studies either reversing dysfunctional microglial states, for example by direct genetic manipulation of key genes, or transplanting homeostatic microglia are needed to identify whether the reversal of microglial dysfunction and enhanced microglial trophic support could slow ALS pathogenesis.

## Conclusions

Overall, the evidence presented in this study demonstrates that microglia change their morphologies prior to immunophenotype changes. We hypothesise these morphological changes aid the disease-related reactivity to neuronal pTDP-43 aggregates, which triggers immunophenotype changes; first, to a phagocytic state then to a dysfunctional one.

## List of abbreviations

ALS: amyotrophic lateral sclerosis
DAM: disease-associated microglia
HLA-DR: human leukocyte antigen – DR isotype
HuBB: Neurological Foundation Human Brain Bank
Iba1: ionised calcium binding adaptor molecule 1
KNN: k-nearest neighbours
pTDP-43: phosphorylated TAR DNA-binding protein 43
rNLS: bigenic *NEFH*-tTA/*tetO*-hTDP-43ΔNLS mice
ROI: region of interest
TDP-43: TAR DNA-binding protein 43
WOD: weeks off doxycycline

## Declarations

## Ethics approval and consent to participate

All human brain tissue was donated to the Neurological Foundation Human Brain Bank following donor and donor family consent, and its use in this study was approved by the University of Auckland Human Participants Ethics committee (protocol number 011654). Animal ethics approval was obtained from The University of Queensland (#QBI/131/18), and experiments were conducted in accordance with the Australian code of practice for the care and use of animals for scientific purposes.

## Consent for publication

Not applicable

## Availability of data and materials

The datasets used and/or analysed during the current study available from the corresponding author on reasonable request.

## Author contributions

MEVS, AKW, and ELS conceptualized the study. RLMF and MAC collected and processed the human tissue. CT carried out pathological assessment of all cases. MEVS and MM carried out the experimental procedures. MEVS and ELS developed the custom image analysis pipelines. MEVS and MM carried out quantification. Data analysis and interpretation were carried out by MEVS, MM, and ELS. Manuscript was prepared by MEVS and ELS. All authors read and approved of the final manuscript.

## Acknowledgements

This publication is dedicated to the incredible patients and families who contribute to our research. We thank Marika Eszes at the Centre for Brain Research, University of Auckland, New Zealand, and the Neurological Foundation of New Zealand for their ongoing financial support of the Human Brain Bank. The imaging data reported in this paper were obtained at the Biomedical Imaging Research Unit, operated by the Faculty of Medical and Health Sciences’ Technical Services at the University of Auckland.

