## Supplementary figures for "Neuronal TDP-43 aggregation drives changes in microglial morphology prior to immunophenotype in amyotrophic lateral sclerosis"

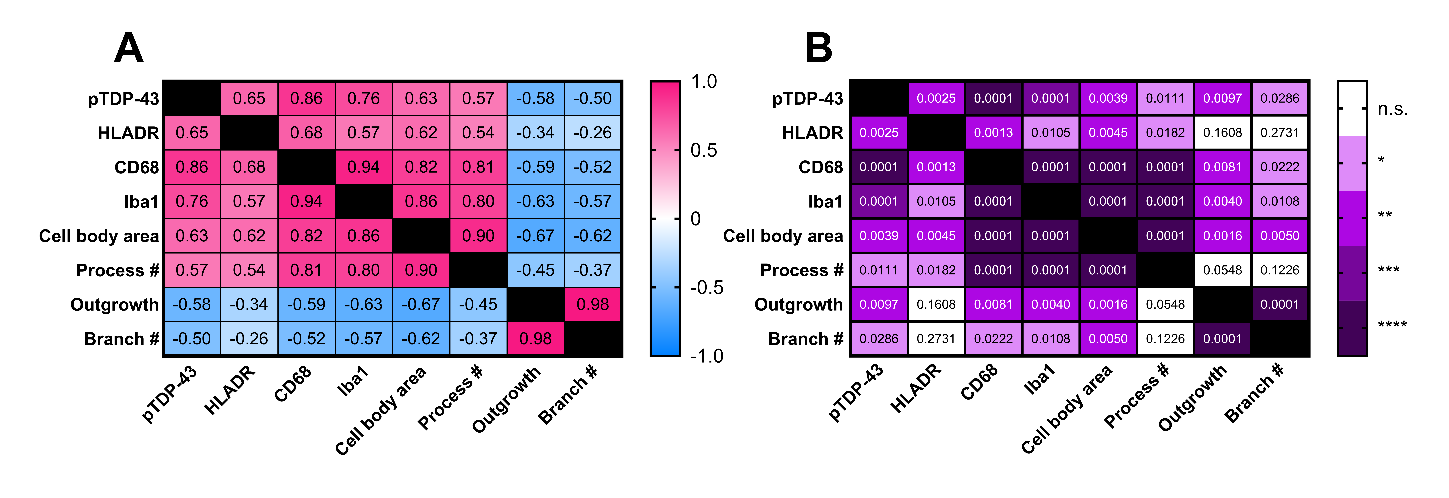

**Figure S1:** **Microglial morphologies correlate with CD68 and Iba1 levels in human ALS.** Measures of pTDP-43 pathology, microglial marker intensities, and microglial morphology were sequentially correlated in all cases (n = 20) using Spearman correlations. The resulting r (**A**) and p (**B**) values from each correlation are presented in the correlation matrices and colour coded relative to strength. When r ≤ -0.7 or r ≥ 0.7 and p ≤ 0.05, correlations were considered statistically significant and strong. When -0.7 < r ≤ -0.4 or 0.7 > r ≥ 0.4 and p ≤ 0.05, correlations were considered statistically significant and moderate. Significance of correlation strengths: *p ≤ 0.05, **p ≤ 0.01, ***p ≤ 0.001, ****p ≤ 0.0001.

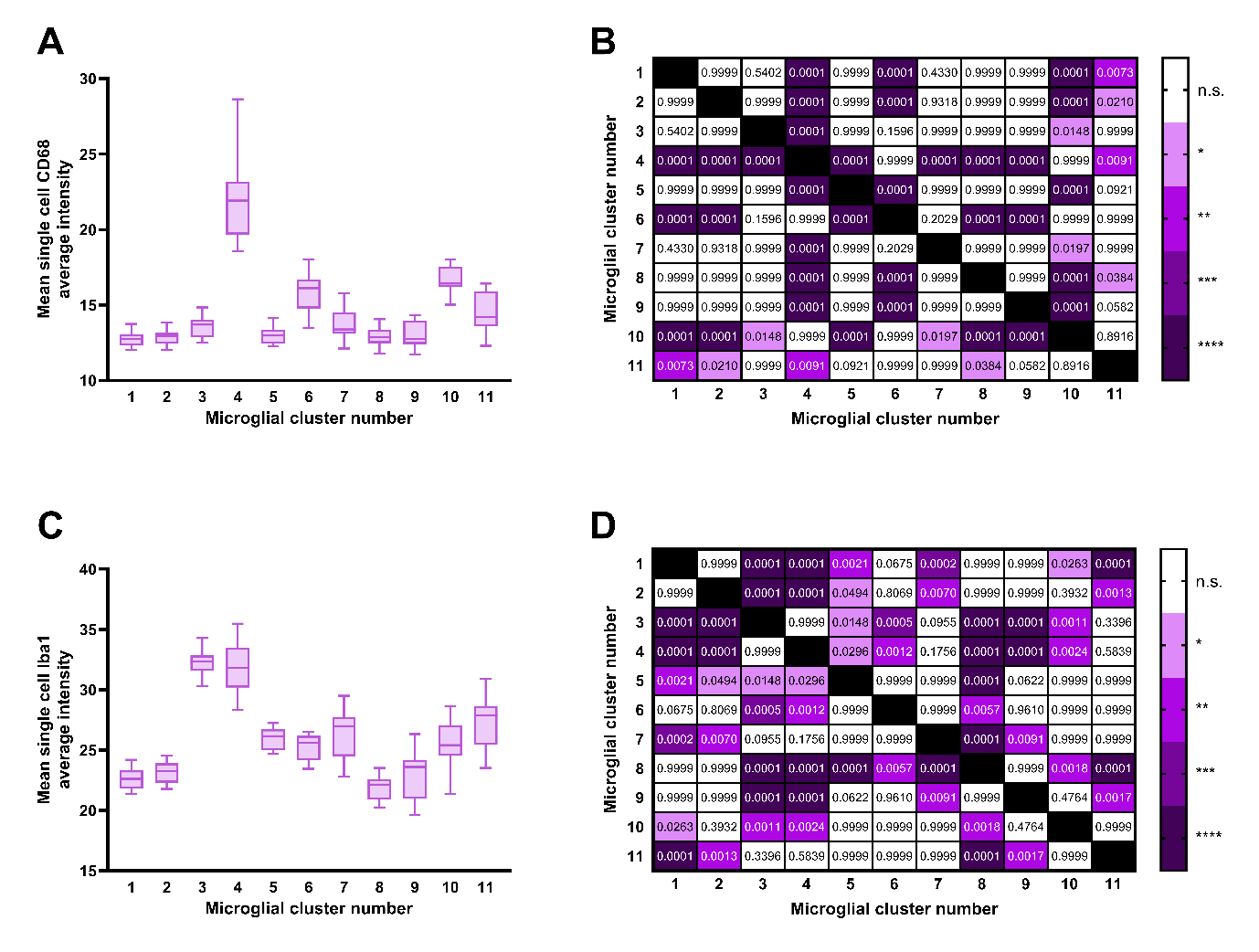

**Figure S2: Microglial clusters enriched in human ALS high Iba1 and CD68 levels.** Louvain clustering was carried out using the single-cell average intensities of CD68, HLA-DR, and Iba1, and morphological measures (cell body area, process number, outgrowth, and branch number) from a randomly subsampled 7,340 microglia per case group (22,020 cells total), resulting in 11 unique clusters. The mean single-cell CD68 and Iba1 average intensity was compared between clusters using a Kruskal-Wallis test with Dunn’s multiple comparisons test (**A** and **C**); data presented as a box and whisker graph, with minimum, maximum, and median shown (n = 20). The significance of mean single-cell CD68 and Iba1 average intensity differences between clusters are shown as heatmaps (**B** and **D**): *p ≤ 0.05, **p ≤ 0.01, ***p ≤ 0.001, ****p ≤ 0.0001.

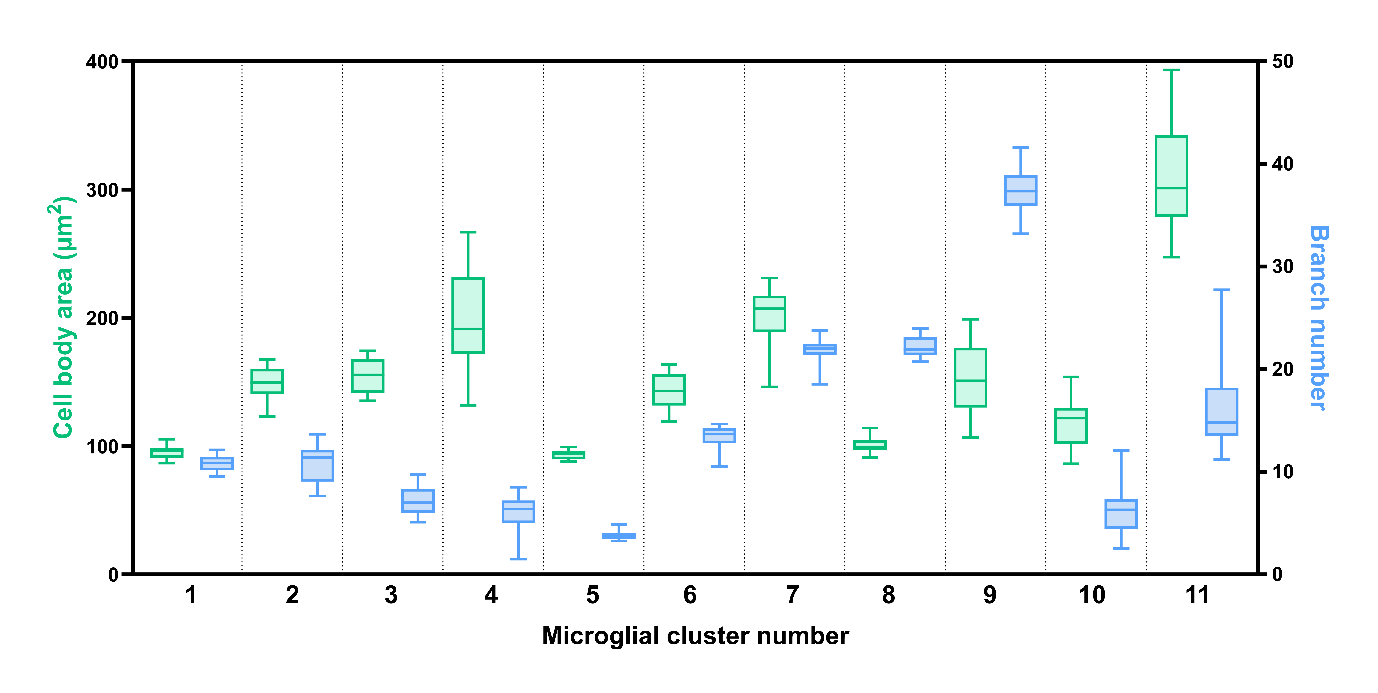

**Figure S3:** **Microglial clusters enriched in human ALS are characterised by high cell body area and low branch number.** The mean cell body area and single-cell branch number were determined for all clusters; data presented for ALS-depleted and ALS-enriched clusters as a box and whisker graph, with minimum, maximum, and median shown (n = 20).
